# Single-cell transcriptomics reveals immune response of intestinal cell types to viral infection

**DOI:** 10.1101/2020.08.19.255893

**Authors:** Sergio Triana, Megan L. Stanifer, Mohammed Shahraz, Markus Mukenhirn, Carmon Kee, Diana Ordoñez-Rueda, Malte Paulsen, Vladimir Benes, Steeve Boulant, Theodore Alexandrov

## Abstract

Human intestinal epithelial cells form a primary barrier protecting us from pathogens, yet only limited knowledge is available about individual contribution of each cell type to mounting an immune response against infection. Here, we developed a pipeline combining single-cell RNA-Seq and highly-multiplex RNA imaging and applied it to human intestinal organoids infected with human astrovirus, a model human enteric virus. We found that interferon controls the infection and that astrovirus infects all major cell types and lineages with a preferential infection of proliferating cells. Intriguingly, each intestinal epithelial cell lineage has a unique basal expression of interferon-stimulated genes and, upon astrovirus infection, undergoes an antiviral transcriptional reprogramming by upregulating distinct sets of interferon-stimulated genes. These findings suggest that in the human intestinal epithelium, each cell lineage plays a unique role in resolving virus infection. Our pipeline can be applicable to other organoids and viruses, opening new avenues to unravel roles of individual cell types in viral pathogenesis.

## Introduction

The small intestine is responsible for most nutrient absorption in humans. It is composed of various cell types each performing specific functions contributing to homeostasis (Peterson & Artis, 2014). The main cell types found in the intestinal epithelium are the absorptive enterocytes, the mucus-secreting goblet cells, the hormone-producing enteroendocrine cells, the antimicrobial peptide secreting Paneth cells and the stem cells. Due to the constant challenges present in the lumen of the gut, intestinal epithelial cells are turned over every five days. This constant self-renewal is organized along the crypt-villus axis and is supported by the stem cells located in the crypts, themselves supported by interlaying Paneth cells (Kretzschmar & Clevers, 2016). Differentiation of the stem cells to the various intestinal cell lineages requires a Notch/Wnt-dependent bifurcation towards either absorptive or secretory progenitor cells. Absorptive progenitor cells give rise to the enterocyte cells while the secretory progenitors differentiate into enteroendocrine, goblet, tuft or Paneth cells (Sancho *et al*, 2015; Gehart & Clevers, 2019). Over the past 10 years, intestinal organoids have been developed and have emerged as the best surrogate model that mimics the differentiation and function of the intestinal epithelium (Sato *et al*, 2011, 2009).

Human intestinal epithelial cells (hIECs) play key roles in protecting us from environmental pathogen- and commensal-related challenges. They act as a first-layer physical barrier of the host defense and mount response upon infection (Martens *et al*, 2018). However, only limited knowledge is available about viral pathogenesis in the intestinal epithelium in the context of the cell types, in particular how different hIEC types contribute to the immune response and clearance of the viral infection. This gap of knowledge is caused by the challenges associated with reproducing the multi-cellular complexity of the human intestinal epithelium. This leads to the current situation where for a majority of enteric viruses, essential questions of viral pathogenesis have been mostly addressed in immortalized cell lines, a model with a limited capacity to reproduce intestinal epithelium in its multi-cellular complexity and missing key phenomena such as cell differentiation. Interestingly, increasing evidence suggests that stem cells are intrinsically resistant to viral infection (Belzile *et al*, 2014; Wolf & Goff, 2009). It was recently shown that pluripotent and multipotent stem cells exhibit intrinsic expression of interferon stimulated genes (ISGs) in an interferon-independent manner. This basal expression of ISGs has been proposed to be responsible for the resistance of stem cells to viral infection (Wu *et al*, 2018). Upon differentiation, stem cells lose expression of these intrinsic ISGs and become responsive to interferon (IFN) (Wu *et al*, 2018). These observations suggest that, at least *in-vitro*, stem cells and differentiated cells use different strategies to fight viral infection. Whether such cell-type-specific antiviral strategies exist *in-vivo* where stem cells differentiate into different tissue-specific lineages remains unknown. Answering these questions requires using physiologically relevant *ex vivo* models as well as single-cell methods able to dissect the heterogeneity of host-pathogen interactions of different cell types and of individual cells within a population.

Here, we established a pipeline to investigate cell-type-specific viral pathogenesis in a tissue-like environment. The pipeline integrates single-cell RNA sequencing and multiplex RNA *in situ* hybridization imaging of enteric virus-infected human ileum-derived organoids. In order to enable cell types-specific analyzes in organoids, we have created a single-cell RNA-Seq atlas of human ileum biopsies at the highest to-date resolution. Applying this pipeline to organoids infected by human enteric virus astrovirus (HAstV1), we have characterized HAstV1-infection of various primary hIEC types and determined that HAstV1 is able to infect all intestinal cell types with a preference for proliferating cells. Single cell transcriptomic analysis revealed a cell-type-specific transcriptional pattern of immune response in human intestinal organoids. We found that both at steady-state and during viral infection, each intestinal cell type has unique expression profiles of interferon-stimulated genes creating a distinct antiviral environment.

## Results

### HAstV1 infects human human intestinal epithelial cells causing IFN-mediated response

To unravel how intestinal epithelium cells respond in a cell-type-specific manner, we investigated human ileum-derived organoids infected with human pathogen astrovirus (HAstV1). Human astrovirus 1 (HAstV1) is a small non-enveloped, positive strand RNA virus which is found worldwide. HAstV1 infections lead to gastroenteritis and can lead to death in children and immunocompromised adults (Cortez *et al*, 2017; Walter & Mitchell, 2003). To confirm the ability of HAstV1 to replicate in intestinal cells, we showed that HAstV1 is fully capable of infecting the transformed hIECs, Caco-2 cells, as can be seen by the increase in the number of infected cells and the increase in the amount of viral genome copy number over time (Fig.S1A-C). Following to the infection with HAstV1, Caco-2 cells mounted an intrinsic immune response characterized by the production of both type I (IFN**β**1) and type III interferons (IFN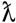) (IFNs) (Fig.S1D). This IFN-mediated response constitutes an antiviral strategy by Caco-2 cells as a pre-treatment of cells with either IFN controlled HAstV1 infection (Fig.S1E-F). This is consistent with previous work reporting that IFN controls HAstV1 infection (Marvin *et al*, 2016; Guix *et al*, 2015). Complementarily, investigating another human intestinal epithelial cell type, T84, which is described to be more immunoresponsive (Stanifer *et al*), we found it to be less infectable by HAstV1 unless the IFN-mediated signaling was suppressed by the loss of both the type I and III IFN receptors (dKO) (Fig.S1G). Together these data confirm the function of IFNs in controlling HAstV1 infection in human intestinal epithelial cells and illustrate the importance of investigating how different cell-types mount an interferon response to counteract viral infection.

### HAstV1 infects human ileum-derived organoids

To investigate whether different cell types in the human intestinal epithelium respond to pathogen challenges by mounting a distinct IFN-mediated response, we exploited human intestinal organoids. Human intestinal organoids are an advanced primary cellular system recapitulating the cellular complexity, organization, and function of the human gut and enabling controlled investigations of enteric infection not feasible in the human tissue (Stanifer *et al*, 2020). Intestinal organoids were prepared from stem cell-containing crypts isolated from human ileum resections (Sato *et al*, 2011). The structural integrity and cellular composition of the organoids were verified by immunofluorescence staining of adherens and tight junctions and markers of various intestinal epithelial cell types (Fig.1A and data not shown) (Pervolaraki *et al*, 2017). These organoids were readily infectable by HAstV1 as can be seen by the detection of infected cells and efficient replication of the HAstV1 genome overtime (Fig.1B-D). Infection of organoids by HAstV1 induces a strong intrinsic immune response characterized by the production of both type I IFN (IFNβ1) and type III interferon (IFNλ) (Fig.1E-F). Interestingly, this response was significantly stronger than the one generated by the transformed hIEC lines (Fig.S1), further highlighting the importance of using primary organoids when characterizing host-pathogen interactions. Pre-treatment of organoids with either type I or type III IFN significantly reduced HAstV1 infection (Fig.1G). This strong antiviral activity of IFN against HAstV1 is consistent with the previous reports using both transformed cell lines and human organoids (Kolawole *et al*, 2019) and with the well-described function of IFNs in controlling enteric virus infection at the intestinal epithelium (Lee & Baldridge, 2017).

### Single-cell RNA-Seq atlas for human ileum biopsies

To characterize the response of intestinal organoids to enteric virus infection at the single-cell level, we exploited single-cell RNA sequencing (scRNA-Seq) by using a 10x Genomics platform (Zheng *et al*, 2017). Over the past years, scRNA-Seq emerged as a major approach to reveal cell types, lineages, and their transcriptional programs in tissues. Yet, applying scRNA-Seq to organoids is still challenging due to the available dissociation protocols optimized predominantly for tissues, as well as due to understudied differences between tissues and organoids. In light of these challenges, prior to analysis of ileum-derived organoids, we performed a comprehensive scRNA-Seq analysis of human ileum biopsies with the aim to create a cell type-annotated reference single-cell atlas of human ileum. For this, human ileum biopsies were subjected to scRNA-Seq followed by data integration and clustering that revealed the presence of multiple cell subpopulations (Fig.2A). We have identified the cell types represented in the subpopulations by finding subpopulation-specific markers and matching them to the known marker genes of the hIEC types. The majority (>87.5%) of the cells isolated from the biopsies corresponded to hIECs while a small fraction (12.4%) corresponded to stroma and immune cells (Fig.2A). Our data are in agreement with the recently published scRNA-Seq data of ileum biopsies (Wang *et al*, 2020). However, our atlas represents more detailed information about cell types containing 14 cell types vs seven cell types in (Wang *et al*, 2020) which is especially important for validating cell types represented in organoids. Additionally, our data contain a larger number of cells (8800 cells vs 6167 cells) (Fig.S2A-F) thus providing a stronger statistical power for finding cell-type-specific markers (Fig.2B and Fig.S3A). For all cell types represented in our scRNA-Seq dataset of human ileum biopsies, we selected the most differentially expressed genes as markers (Fig.2B). The pseudo-time analysis (Street *et al*, 2018) confirmed the expected bifurcation of differentiation of stem cells along two distinct trajectories toward either enterocytes (absorptive function) or goblet cells (secretory function) (Fig.2C). Each of these two lineages are characterized by specific gradients and waves of gene expression along the differentiation trajectories (Fig.2D). Together these data provide detailed single-cell atlas of intestinal epithelial cell types and their transcriptional programs in the human ileum.

### Human ileum-derived organoids reproduce tissue cell types and lineages

Human ileum organoids were subjected to scRNA-Seq followed by determining the identities of the detected cell types by using cell-type-specific marker genes from the ileum biopsies (Fig. 3A). Most of the anticipated and observed cell types in biopsies were also found in organoids with the exception of Paneth and tuft cells (Fig.3A). This is consistent with previous reports describing the absence of these two intestinal types in organoids (Fujii *et al*, 2018) and with the fact that some cell types require specific differentiation protocols to be present in organoids (Wu *et al*, 2018; Rouch *et al*, 2016; Ding *et al*, 2020). Correlation of class-average transcriptional profiles between organoids and biopsies revealed an overall high correlation within every cell type with the Pearson r≥0.75 for stem cells and transit-amplifying cells; r≥0.6 for enterocytes, immature enterocytes and enterocytes progenitors; and r≥0.5 for goblet and secretory transit-amplifying cells (Fig.S4). The high correlation for stem cells, transit-amplifying cells, and enterocytes in organoids to their *in vivo* counterparts (Fig.S4C-D), despite the expected differences in the exposure of *in vivo* enterocytes to the gut luminal microenvironment, further highlights the ability of organoids to faithfully recapitulate tissue transcriptional programs.

**Figure 1.**
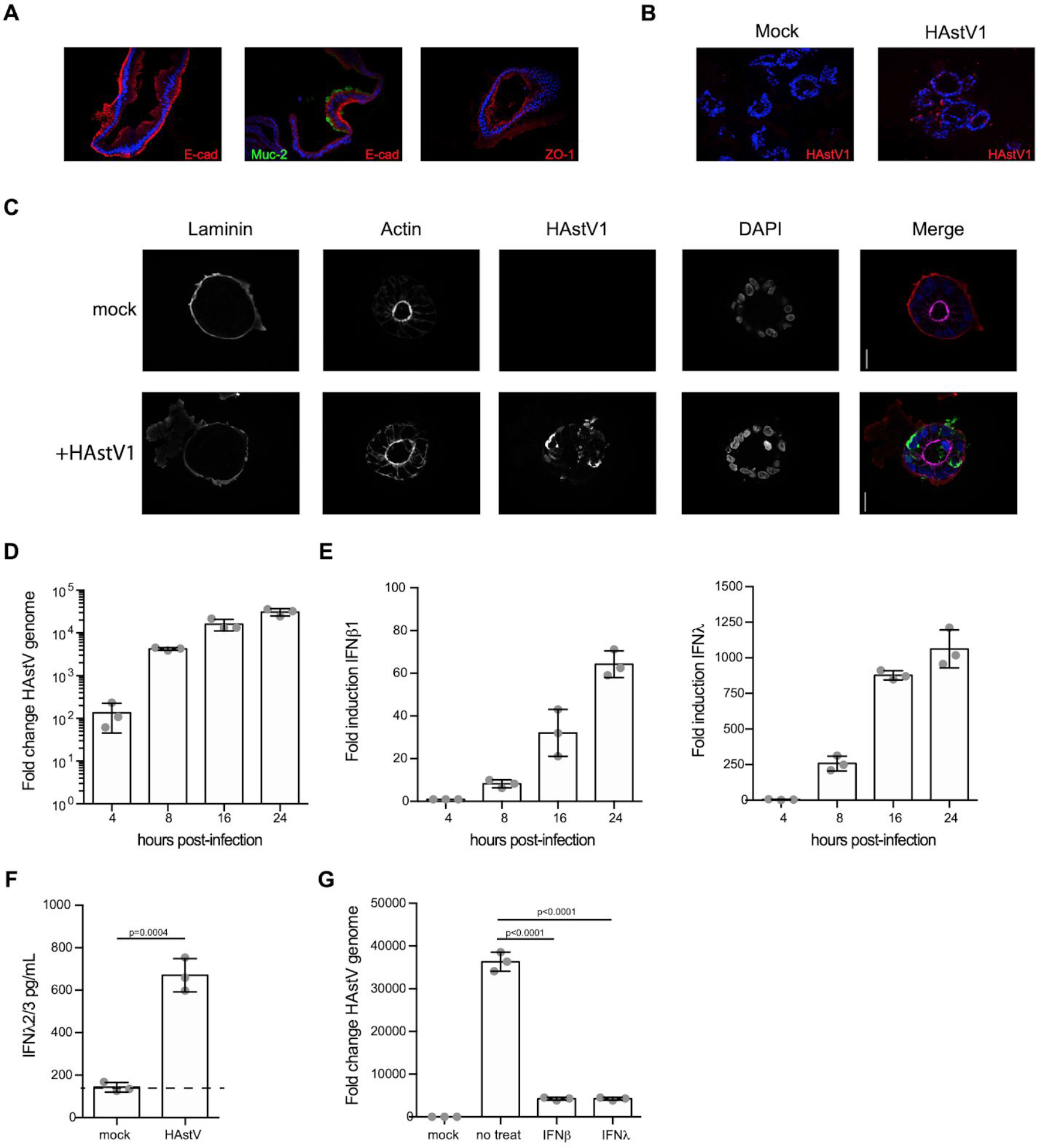
Interferons protect human intestinal organoids from HAstV1 infection. **A**. Cryo-sections of human organoids were analyzed for the presence of enterocytes (E-cad), Goblet cells (Muc-2) and tight junctions (ZO-1) by indirect immunofluorescence. Nuclei are stained with DAPI. **B**. Human intestinal organoids were incubated with media (mock) or infected with HAstV1. 16hpi organoids were frozen, cryo-sectioned, and HAstV1 infected cells were visualized by indirect immunofluorescence (HAstV1 (red), nuclei (DAPI, blue). **C**. Human intestinal organoids were incubated with media (mock) or infected with HAstV1. 16hpi organoids were fixed and the presence of HAstV1 infected cells (green) were visualized by indirect immunofluorescence. Apical and basolateral membranes were immunostained for actin (magenta) and Laminin (red) respectively. Nuclei are stained with DAPI (blue). **D**. Human intestinal organoids were infected with HAstV1. At indicated time post-infection the increase in viral copy number was determined by q-RT-PCR. **E**. Same as D but the induction of type I (IFN1) and III (IFNA) IFN were evaluated. **F**. Human intestinal organoids were incubated with media (mock) or infected with HAstV1. 24hpi the presence of IFNA in the media was tested by ELISA. Dotted line indicates detection limit of the assay. **G**. Human intestinal organoids were pre-treated for 24h with 2000IU/mL of IFNβ1 or 300ng/mL of IFNλ1-3. Interferons were maintained during the course of infection and the amount of HAstV1 copy numbers was assayed 24hpi by q-RT-PCR. A-G Three biological replicates were performed for each experiment. Representative immunofluorescence images are shown. Error bars indicate the standard deviation. Statistics are from unpaired t-test.

**Figure 2.**
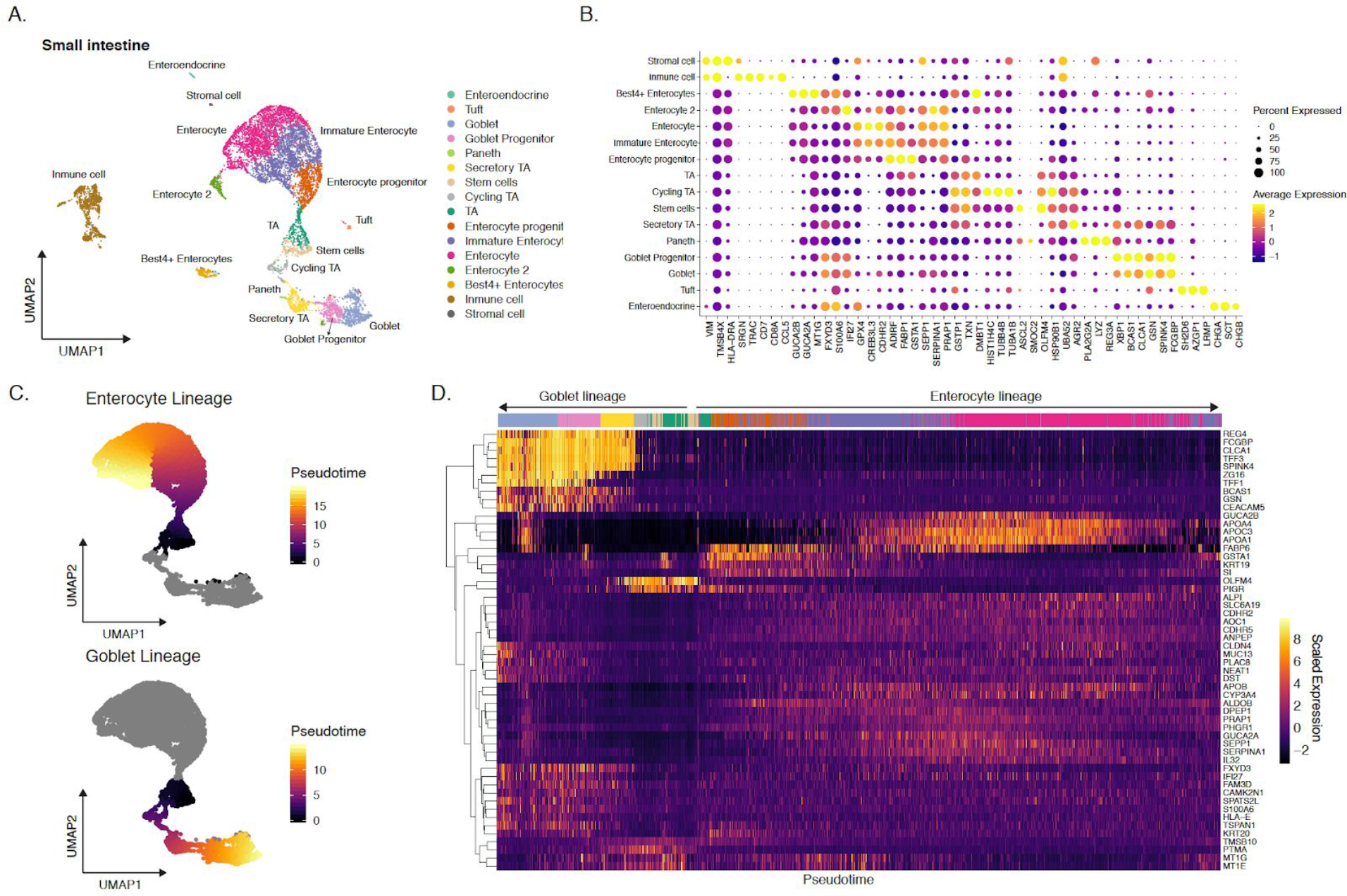
Single-cell profiling of human ileum biopsies. **A**. Uniform manifold approximation and projection (UMAP) embedding of single-cell RNA-Seq data from human ileum biopsies colored by the cell type (n=8,800 cells). **B**. Dot plot of the top three marker genes for each cell type. The dot size represents the percentage of cells expressing the gene; the color represents the average expression across the cell type. Fig.S3A shows an extended list of the marker genes. All cell-type-defining marker genes are provided in Table S2. **C**. Predicted pseudo-time trajectories for the goblet and enterocyte lineages projected onto the UMAP embedding. **D**. Heatmap showing changes in gene expression levels along pseudotime for the enterocytes and goblet lineages. Dendrograms on the left of the heatmap indicate the results of hierarchical clustering of the genes. Colored bars above the heatmap indicate the types of cells ordered along the pseudotime, colored according to A.

**Figure 3.**
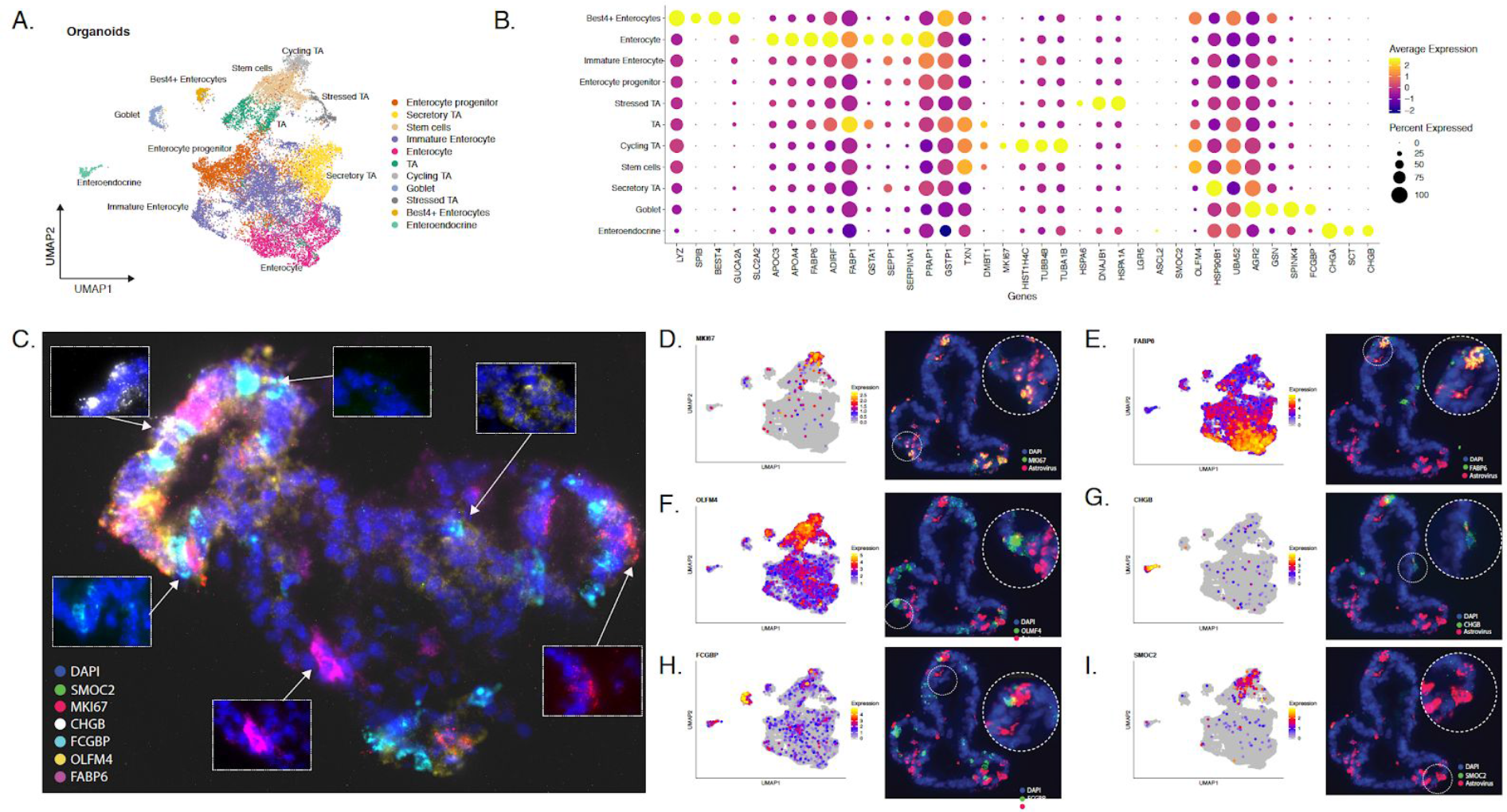
Single-cell profiling and multiplex *in situ* RNA hybridization of human ileum-derived organoids. **A**. UMAP of scRNA-Seq data from human ileum-derived organoids (n=16,682 cells); dots corresponding to cells are colored by the cell type. **B**. Dot plot of the top-three marker genes for each cell type. The dot size represents the percentage of cells expressing the gene; the color represents average expression across the cell type. Fig.S3B shows an extended list of the marker genes. All cell-type-defining marker genes are provided in Table S3. **C**. Representative images showing multiplex *in situ* RNA hybridization of marker genes in an organoid. DAPI in blue. **D-I**. Visualizations for cell-type markers for the major intestinal cell types, showing single-cell expression intensities of the cell-type markers on the UMAP (left) and multiplex *in situ* RNA hybridization; red corresponds to viral RNA and a green corresponds to cell type markers(right).

**Figure 4.**
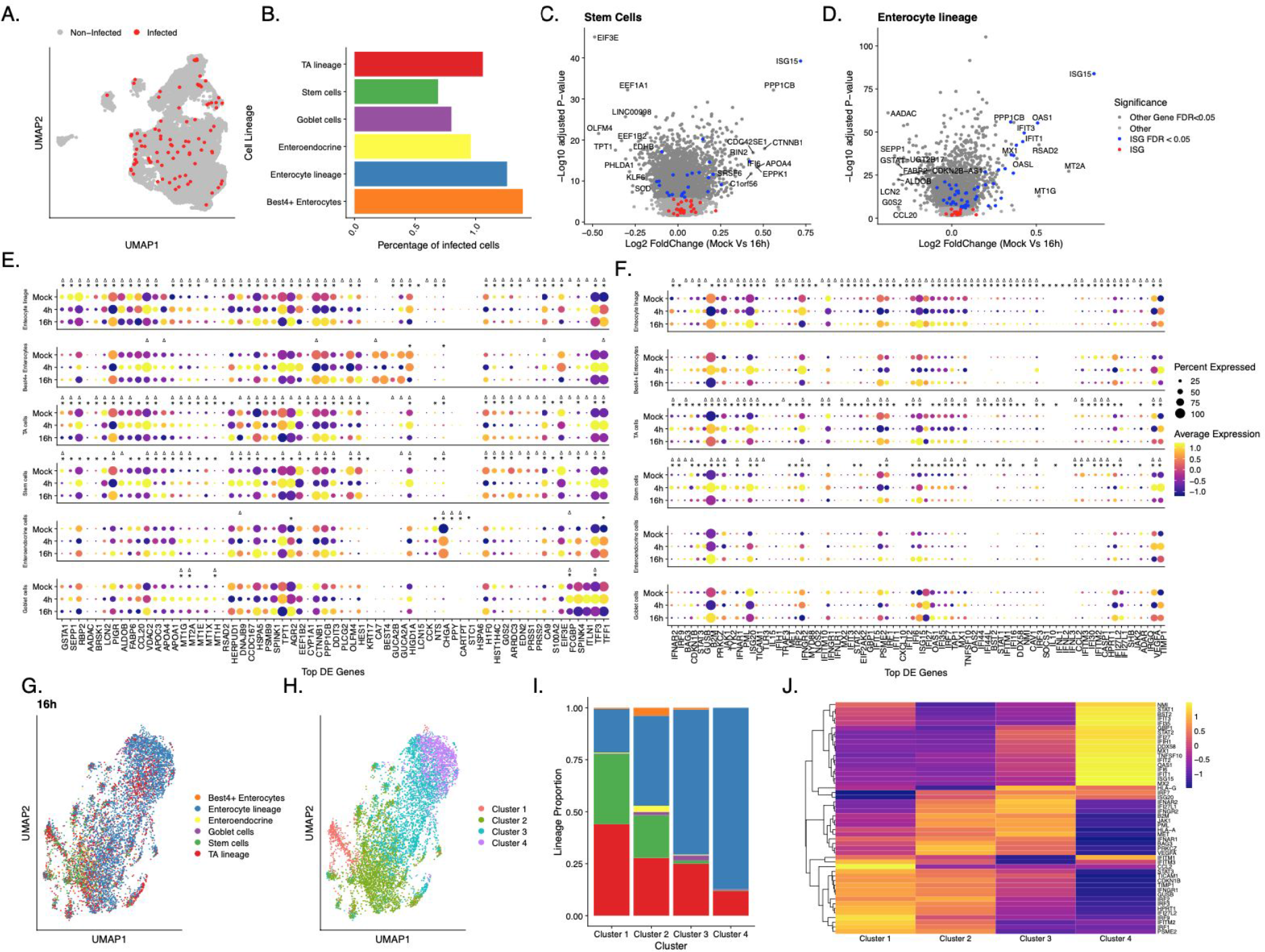
Cell-type-specific ISG induction upon HAstV1 infection of human ileum organoids. **A**. UMAP embedding of human ileum-derived organoids highlighting HAstV-infected cells (cells where astrovirus transcripts were detected in scRNAseq). **B**. Percentage of cells infected with astrovirus detected in each cell lineage and type. **C-D**. Volcano plots of genes that are differentially expressed in mock relative to 16hpi, showing the statistical significance (-log10 adjusted p-value) vs log2 fold change (mock/16hpi) for the stem cells (C) and enterocyte lineage (D). **E**. Dot plot of top lineage-specific expression changes upon HAstV1 infection for mock, 4hpi and 16hpi organoids.. **F**. Same as in **E** but for ISG. The dot size represents the percentage of cells expressing the gene; the color represents the average expression across the lineage. Triangle and stars represent significantly changing genes (FDR < 0.05) in Mock vs 4hp.i and Mock vs 16p.i respectively **G**. UMAP embedding of scRNA-Seq data from human ileum-derived organoids based on the significantly changing ISGs for cells at 16hpi. **H**. Unsupervised clustering of the UMAP data from G. **I**. Distribution of cell lineages and types in the clusters from H. **J**. Heatmap of differentially expressed ISGs across the clusters from H.

### Multiplex *in situ* RNA hybridization visualizes multi-cellular organization and infection of organoids

ScRNA-Seq is a powerful and robust tool to investigate cell-cell heterogeneity, outline cell types, and determine their marker genes. However, it lacks the capacity for spatial mapping of the discovered information. To overcome this limitation, we employed the HiPlex RNAscope method for multiplexed RNA *in-situ* hybridization (Hashikawa *et al*, 2020). This method allows for highly-sensitive single-molecule detection of 12 different transcripts, with four transcripts detected simultaneously in each round of hybridization (Fig.S5A). To determine the best cell type-specific marker genes expressed in organoids, we have repeated the differential analysis and identified marker genes for each individual intestinal cell type; see top markers in Fig.3B and extended list in Fig.S3B. From this list, we selected 11 genes for the multiplexed RNA *in-situ* hybridization which were either uniquely expressed in a specific cell type or had less background expression among cells of other types (Fig.S5B). The probes directed against SMOC2, MKI67, CHGB, FCGBP, OLFM4, FABP6 and LYZ showed discrete staining overlapping within cell borders suggesting cell type specificity. On the other hand, probes directed against SLC2A2, BEST4, LGR5 and SPIB showed either no signal or a background unspecific signal (Fig.S5B). RNA imaging of 11 cell type marker genes, previously not applied at such a high level of multiplexity in organoids (Tsai *et al*, 2017; Navis *et al*, 2019; Seino *et* al, 2018; Smillie *et al*, 2019), allowed us to validate the expression of these genes and provided a spatial insight into the cell organization in particular with respect to proliferating areas within an organoid (Fig.3C).

To identify which cell types were infected by HAstV1, we designed an HAstV1 RNA-specific probe and performed the multiplexed RNA *in situ* hybridization on infected organoids. (Fig.3D-I and Fig.S5B). *In situ* hybridization of mock infected organoids using the HAstV1 RNA-specific probe revealed no labelling thus validating the specificity of the HAstV1 probe. In HAstV1-infected organoids, viral RNA was strongly co-localized with the expression of MKI67 (Fig.3D), a marker of proliferation which in our organoid system is expressed predominantly in cycling transit-amplifying cells (Fig.3B). Additionally, mature enterocytes (marker FABP6) were also found to be infected by HAstV1 (Fig.3E). Viral RNA was also detected, although to lesser extent, in stem-like cells (marker OLFM4) (Fig.3F), enteroendocrine cells (marker CHGB) (Fig.3G), as well as goblet cells (marker FCGBP) (Fig.3H). These results indicate the potential of HAstV1 to infect most hIECs and are in agreement with a recent report showing that HAstV1 can infect most human intestinal epithelial cell types (Kolawole *et al*, 2019).

### HAstV1 evokes cell-type-specific transcriptional response

To address whether the cell types present in the intestinal organoids differently respond to viral infection, we performed scRNA-Seq of HAstV1-infected organoids. Single-cell transcriptional analysis confirmed that all intestinal epithelial cell types could be infected by HAstV1 (Fig.4A-B). Differential gene expression analysis of mock-infected organoids and organoids at 4 and 16 hours post-infection with HAstV1 revealed upregulation of multiple pathways upon infection, including the pathways linked to infection, pro-inflammatory response and interferon mediated response (Fig.S6). Interestingly, differential gene expression analysis integrating the different cell types revealed a cell-type-specific transcriptional response upon virus infection (Fig.4C-E). This response includes canonical marker genes for the detected cell types *e*.*g*. upregulation of APOA1 in enterocytes, downregulation of OLFM4 in stem cells and TA cells, and upregulation of CHGA in enteroendocrine cells (Fig.4E). Most importantly, our analysis revealed that different ISGs are upregulated in different cell types upon viral infection (Fig.4C-D and Fig.S6). These findings suggest that each cell type present in our intestinal organoids has a unique transcriptional program to combat viral infection.

To better evaluate the involvement of individual cell types into the interferon-mediated response to HAstV1 infection (Fig.1), we performed detailed differential analysis using ISGs only (Fig.4F). For the enterocytes lineage, 68 out of 80 ISGs were found to significantly change either at 4hpi or 16hpi compared to mock with almost all significant ISGs upregulated at 16hpi. Other cell types including stem and TA cells demonstrated their cell-type-specific changes at both early (4hpi) and late (16hpi) time points (Fig.4F).

Surprisingly, the unsupervised mapping of all cells based on their ISG profiles revealed an organization of cells according to the cellular lineage (Fig.4G and Fig.S7B-C). Further clustering of cells based on their ISG profiles revealed four clusters (Fig.4H and Fig.S7E-F) with cluster 1 being mostly composed of stem and TA cells and cluster 4 being mostly composed of enterocytes (Fig.4I and Fig.S7H-I). Differential analysis revealed distinct patterns of ISGs expressed upon HAstV1 infection which are specific to the cell types represented in the outlined clusters (Fig.4J and Fig.S7K-L). Importantly, similar analyses of mock-infected organoids also revealed a lineage-specific clustering of cells according to their ISG expression profiles (Fig.S7A,D,G and J). Even in mock-infected organoids each cluster expresses different basal levels of distinct ISGs (Fig.S7J). Altogether, these findings strongly support that each intestinal cell lineage is characterized by a distinct basal signature of ISG expression and that upon viral infection, each intestinal cell lineage responds differently by mounting a distinct immune response characterized by cell-type-specific patterns of ISG expression.

## Discussion

In this work we exploited human intestinal organoids, scRNA-Seq, and multiplex RNA *in situ* hybridization to characterize the response to infection by the enteric virus HAstV1 in a primary model system that recapitulates the cellular complexity of the human gut. Using multiplex RNA *in situ* hybridization spatially correlating viral infection to cell types of an organoid, we found that HAstV1 infects all detected cell types but displays a preference for proliferating cells. ScRNA-Seq confirmed that HAstV1 could infect all intestinal cell types. Importantly, it revealed that each individual intestinal cell lineage mounted a distinct transcriptional response upon viral infection. We found cell-type-specific transcriptional patterns of ISGs expression with different lineages characterized by the expression of unique ISGs. Importantly, our analysis also revealed that the basal level of ISG expression in non-infected organoids was also cell-type-specific. These findings strongly suggest that within the intestinal epithelium, each individual cell type utilizes a different strategy to combat viral infection and has a different function in the clearance of the pathogen.

Investigating host-pathogen interactions and viral pathogenesis in tissue is challenging. It requires identifying cell types composing the tissue and correlating virus infection to each cell type within its spatial context. Here, we have developed a technological pipeline combining scRNA-Seq and multiplex RNA *in situ* hybridization to overcome these limitations. Associating transcriptional changes to specific cell types in scRNA-Seq data requires validated cell-type-specific marker genes. Despite the availability of single-cell atlases for other parts of the intestine published recently (Smillie *et al*, 2019), no reference atlas was available for the small intestine when we started this work. Recently, Wang *et al* published a scRNA-Seq dataset of human ileum with a total of 6167 epithelial cells and 7 cell types annotated (Wang *et al*, 2020). Our scRNA-Seq dataset from human ileum biopsies is in line with the data published by (Wang *et al*, 2020) however contains more cells (of 8800 vs 6167) and twice more cell types annotated (14 vs 7). Having a comprehensive coverage of cell types and differentiation lineages in a reference atlas is especially important when working with organoids due to possible transcriptional changes between organoid and tissue cells as well as the presence of not fully-differentiated cells that otherwise may lead to false annotations. Here, we have validated that the transcriptional profiles in cells of the major epithelial cell types in the organoids correlate highly to the profiles of their *in vivo* counterparts.

Applying scRNA-Seq to study infection in multi-cellular models or tissue is still challenging as it is a sophisticated technology requiring optimization of cell dissociation, fixation and pathogen inactivation. This leads to the current situation when scRNA-Seq is applied mainly to study viral infection in cell lines (Wyler *et al*, 2019; O’Neal *et al*, 2019; Cristobal Vera *et al*, 2019) or primary tissue where one cell type is present or only mature cells are present like PBMCs (Zanini *et al*, 2018) or lung epithelial cells (Cao *et al*, 2020). Optimizing our pipeline, we have evaluated whether fixation or sorting should be performed and concluded that using live cells without fixation provided the best quality scRNA-Seq data for infected organoids and that using sorting introduces no significant differences (Fig.S8). Overall, we collected a to-date largest scRNA-Seq dataset from human intestinal organoids with 16,682 cells whereas previous studies reported on ∼6000 cells (Fujii *et al*, 2018) and ∼2000 cells (Chen *et al*, 2019). This is particularly important for annotating comparatively rare cell types such as cycling or stressed TA cell cells and goblet cells as well as for investigating possible heterogeneity upon infection.

Although scRNA-Seq provides a comprehensive view of transcriptional programs and serves as an excellent tool for outlining cell types, subtypes and differentiation lineages, we found RNA *in situ* hybridization to be crucially important as an orthogonal way to provide spatial information about the expression of marker genes. Using *in situ* hybridization for the viral RNA represents a native approach to integrate information about cell types and infection. First, it can be more sensitive and specific for virus localization compared to immunofluorescence and importantly it represents a fast alternative in case there is no specific antibody available. Second, this approach allowed us to bypass the main limitation of correlating scRNA-Seq with fluorescence microscopy which is often limited by the availability of antibodies for the different cell type markers. Most importantly, using scRNA-Seq and RNA *in situ* hybridization together for cell-type-analysis leads to further advantages compared to complementing scRNA-Seq with detecting cell types using antibodies. A limited correlation between abundances of RNA molecules and cognate proteins can lead to discrepancies of the cell types definitions and/or corresponding cell-type-markers between the transcript and proteins levels. Localizing viral RNA together with cell-type-specific marker genes requires multiplexing. Here, we show the first application of highly multiplex (>10 channels) RNA *in situ* hybridization to infected organoids whereas previous studies reported detection of one or two cell types (Tsai *et al*, 2017; Navis *et al*, 2019; Seino *et al*, 2018; Smillie *et al*, 2019).

Understanding enteric virus pathogenesis and how the intestinal epithelium combats viral infection has become a growing field of interest due to the socioeconomic impact of viral gastroenteritis (Bányai *et al*, 2018). Interferons and the associated interferon-stimulated genes are well known to contribute to the first-line defence against viruses. Among the different types of IFNs, type III IFN (*i*.*e*. IFN**λ**) (Kotenko *et al*, 2003; Sheppard *et al*, 2003) has been shown to be the key antiviral cytokine controlling enteric viruses (Pervolaraki *et al*, 2017, 2018; Pott *et al*, 2011; Nice *et al*, 2015). Our scRNA-Seq analysis confirm that most HAstV1 infected cells show high expression of IFN**λ** and ISGs. On the contrary, type I IFN (IFNB1) was not detected in our scRNA-Seq data, confirming the type III IFN preference of IECs in response to viral infection (Mahlakõiv *et al*, 2015).

Stem cells have been known to be highly resistant to viral infection compared to differentiated cells. It has been shown that although these cells are refractory to IFNs, they express a basal level of ISGs which are responsible for their intrinsic resistance to viral infection (Wu *et al*, 2018). Interestingly, different tissue stem cells show a distinct subset of ISG expression patterns. Differentiation of human embryonic stem cells to endo-, meso- and ectoderm also show that the three different lineages have a different basal level of ISGs (Wu *et al*, 2018). Here, by using intestinal organoids, we could show that in a naturally-differentiating system all intestinal lineages and their progenitor cells display distinct basal expression levels of ISGs. This may contribute to viral tropism by creating an antiviral state restricting virus in a cell-type-specific manner.

The use of intestinal organoids to study host-enteric pathogen interaction is only at its premise (Dutta & Clevers, 2017). Bulk transcriptional analysis of viral infection in intestinal organoids, which are fully immuno-responsive compared to many intestinal-derived cell lines, shows that they induce a typical IFN-mediated antiviral response (Kolawole *et al*, 2019; Lamers *et al*, 2020). ScRNA-Seq of both primary and immortalized virus-infected cell lines revealed a population heterogeneity of both viral replication and the transcriptional response of cells to viral infection (Combe *et al*, 2015; Russell *et al*; Wyler *et al*, 2019). Here, we report the first scRNA-Seq study of viral infection of organoids. Our data not only show an IFN-mediated antiviral response but also highlight how individual intestinal lineages contribute to mounting the overall response by activating unique transcriptional programs characterized by cell-type-specific subsets of ISGs (Fig.4 and S7). Interestingly, we also observed a heterogeneity of the transcriptional response upon infection within cells of the same lineage. However and most importantly, clustering cells based on their ISGs profiles reveals that the differences of the ISG expression between intestinal cell lineages is higher than the heterogeneity between cells of the same lineage (Fig.S7). This cell-type-specificity of expression of ISGs observed for the enterocyte lineage, stem cells, transit-amplifying cells, and goblet cells confirms the potential of using bioengineered tissue-like systems to dissect the cell type-specific immune response upon host-pathogen interactions. From a biological point of view, our findings strongly suggest that individual cell types within the intestinal epithelium exert different functions during pathogenesis and contribute differently to mounting the immune response and to the overall clearance of the pathogen.

We anticipate our pipeline for integrative single-cell RNA sequencing and RNA imaging of organoids to be widely applicable to other organoid models and to any RNA virus with poly(A) tails thus representing a novel approach to study viral pathogenesis and able to provide novel insights into immune response and cell-type-specific transcriptional reprogramming upon infection.

## Materials and Methods

### Cells and viruses

T84 human colon carcinoma cells (ATCC CCL-248) were maintained in a 50:50 mixture of Dulbecco’s modified Eagle’s medium (DMEM) and F12 (GibCo) supplemented with 10% fetal bovine serum and 1% penicillin/streptomycin (Gibco). Caco-2 human colorectal adenocarcinoma (ATCC HTB-37) were maintained in Dulbecco’s modified Eagle’s medium (DMEM) (GibCo) supplemented with 10% fetal bovine serum and 1% penicillin/streptomycin (Gibco). Human Astrovirus 1 (HAstV1) was a kind gift from Stacy Schultz-Cherry, St. Jude Children’s Research Hospital, TN, USA. HAstV1 was amplified in Caco-2 cells and virus-containing supernatants were used for infection experiments.

### Human organoid cultures

Human tissue was received from colon and small intestine resection from the University Hospital Heidelberg. This study was carried out in accordance with the recommendations of the University hospital Heidelberg with written informed consent from all subjects in accordance with the Declaration of Helsinki. All samples were received and maintained in an anonymized manner. The protocol was approved by the “Ethics commission of the University Hospital Heidelberg” under the protocol S-443/2017. Stem cells containing crypts were isolated following 2 mM EDTA dissociation of tissue samples for 1 h at 4°C. Crypts were spun and washed in ice cold PBS. Fractions enriched in crypts were filtered with 70 µM filters and the fractions were observed under a light microscope. Fractions containing the highest number of crypts were pooled and spun again. The supernatant was removed, and crypts were re-suspended in Matrigel. Crypts were passaged and maintained in basal and differentiation culture media (see Table 1).

**Table 1.** Organoids culturing media.

| *Compound* | *Final concentration* |
| --- | --- |
| **Basal media** |  |
| Ad DMEM/F12<br>+GlutaMAX<br>+HEPES<br>+P/S |  |
| Wnt3A | 50% by volume |
| B27 | 1:50 |
| N2 | 1:100 |
| N-acetyl-cysteine | 1mM |
| R-spondin | 10% by volume |
| Noggin | 100ng/mL |
| EGF | 50ng/mL |
| Gastrin | 10mM |
| Nicotinamide | 10mM |
| A83-01 | 500nM |
| Sb202190 | 10uM |
| **Differentiation Media** |  |
| Ad DMEM/F12<br>+GlutaMAX<br>+HEPES<br>+P/S |  |
| B27 | 1:50 |
| N2 | 1:100 |
| N-acetyl-cysteine | 1mM |
| R-spondin | 5% by volume |
| Noggin | 50ng/mL |
| EGF | 50ng/mL |
| Gastrin | 10mM |
| A83-01 | 500nM |
| Sb202190 | 10uM |

### Antibodies and inhibitors

ZO-1 (Santa Cruz Biotechnology) was used at 1/100 for immunostaining; Phalloidin-AF647 (Molecular Probes) was used at 1/200 for immunostaining; Laminin (Abcam) was used at 1/100 for immunostaining, mouse monoclonal against E-cadherin (BD Transductions), and rabbit polyclonal anti-Mucin-2 (Santa Cruz Biotechnology) were used at 1:500 for immunofluorescence and HAstv1 capsid antibody (Abcam) was used at 1/300 for immunostaining. Secondary antibodies were conjugated with AF488 (Molecular Probes), AF568 (Molecular Probes), and AF647 (Molecular Probes) directed against the animal source.

### RNA isolation, cDNA, and qPCR

RNA was harvested from cells using NuceloSpin RNA extraction kit (Machery-Nagel) as per manufacturer’s instructions. cDNA was made using iSCRIPT reverse transcriptase (BioRad) from 250 ng of total RNA as per manufacturer’s instructions. q-RT-PCR was performed using iTaq SYBR green (BioRad) as per manufacturer’s instructions, HPRT1 was used as normalizing genes. Primer used:

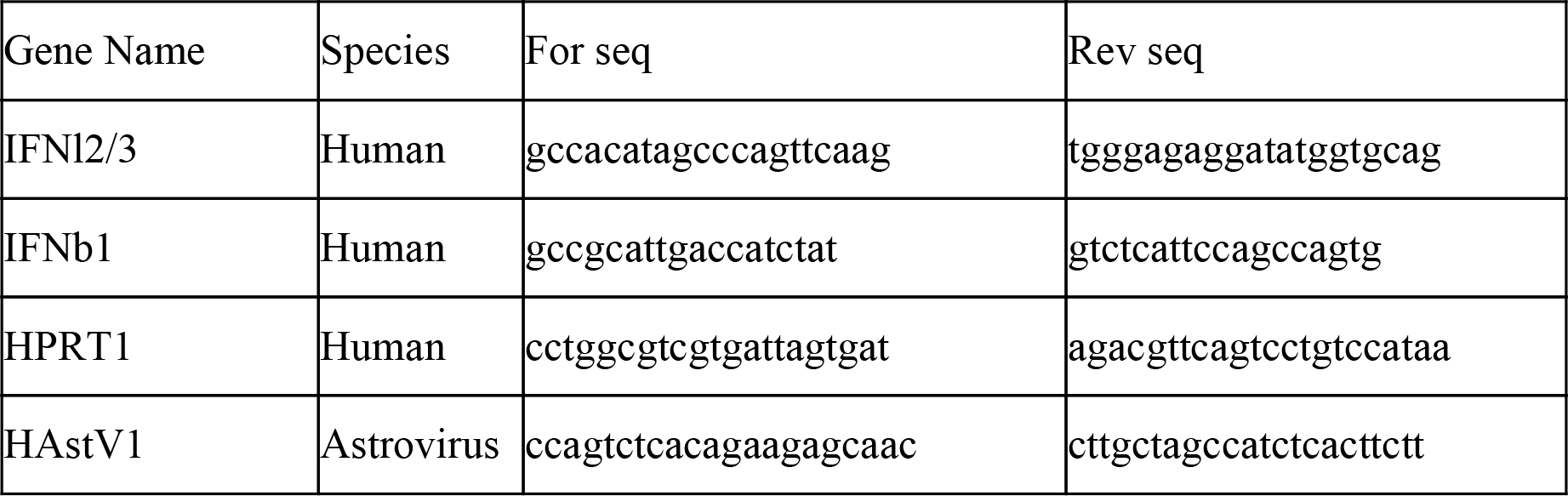

### Indirect immunofluorescence assay

Cells seeded on iBIDI glass bottom 8-well chamber slides. At indicated times post-infection, cells were fixed in 4% paraformaldehyde (PFA) for 20 mins at room temperature (RT). Cells were washed and permeabilized in 0.5% Triton-X for 15 mins at RT. Primary antibodies were diluted in phosphate-buffered saline (PBS) and incubated for 1h at RT. Cells were washed in 1X PBS three times and incubated with secondary antibodies and DAPI for 45 mins at RT. Cells were washed in 1X PBS three times and maintained in PBS. Cells were imaged by epifluorescence on a Nikon Eclipse Ti-S (Nikon).

Organoids were fixed in 2% paraformaldehyde (PFA) for 20 mins at room temperature (RT). Cells were washed and permeabilized in 0.5% Triton-X for 15 mins at RT. Primary antibodies were diluted in phosphate-buffered saline (PBS) and incubated for 1h at RT. Antibody was removed and samples were washed in 1X PBS three times and incubated with secondary antibodies for 45 mins at RT. Antibody was removed and samples were washed in 1X PBS three times and kept in water for imaging. Organoids were imaged on an inverted spinning disc confocal microscope (Nikon, PerkinElmer) with 60x (1.2 numerical aperture, PlanApo, Nikon) water immersion objective and an EMCCD camera (Hamamatsu C9100-23B).

### ELISA

Supernatants were collected at 16 hours post-infection. Supernatants were kept undiluted. IFNλ2/3 was evaluated using the IFNλ2/3 DIY ELISA (PBL Interferon source) as per manufacturer’s instructions.

### Organoid infection

Organoids were moved to differentiation media (Table 1) three days prior to infection. At the time of infection, medium was removed and organoids were resuspended in cold PBS to remove any excess of Matrigel. To allow for infection from both the apical and basolateral sides, organoids were disrupted with a 27G needle 10 times. Mock infection cells were incubated with media and infected samples were incubated with supernatant containing virus. Infections were allowed to proceed for one hour. Cells were then spun down and virus was removed. Cells were washed one time with DMEM/F12 and then resuspended in differentiation media for the course of the experiment.

### Tissue and organoid dissociation for scRNA-seq

Obtained fresh human ileal biopsies were kept in PBS at 4oC until processing using the previously described method (Smillie *et al*, 2019). The biopsy material was cut into small pieces using scissors. The dissociated tissue was then incubated in dissociation solution (HBSS Ca/Mg-Free, 10 mM EDTA, 100 U/ml penicillin, 100mg/mL streptomycin, 10 mM HEPES, and 2% FCS) with 200mL of 0.5M EDTA added at the time of dissociation. Tissue was incubated in dissociation solution for 15mins at 37oC with rotating. Cells were then incubated on ice for 10 mins and then shaken 15-20 times. At this time microscopic examination revealed that crypts had been released and therefore the large pieces of tissue were removed. The crypts were then spun down at 500xg for 5 mins. The supernatant was removed and crypts were washed one time in cold PBS. Crypts were then digested to single cells by incubation with TrpLE for 5-10 mins at 37oC or until microscopic examination showed single cells were present. Cells were then spun at 500xg for 5 mins and washed one time with PBS. Any red blood cells contamination was removed by a 3 min incubation on ice with ACK lysis buffer (Thermo). Cells were then spun at 500xg for 5 mins and washed one time with PBS. Supernatant was removed and the cell pellet was resuspended in PBS supplemented with 0.04% BSA and passed through a 40-µm cell strainer and used directly for single cell RNASeq.

For organoid samples, cultured organoids harvested after 0 (mock), 4 and 16 hours post-infection were washed in cold PBS to remove excess Matrigel and incubated in TrypLE Express for 25 min at 37°C. When microscopic examination revealed that cells had reached a single cell state, they were resuspended in DMEM/F12 and spun at 500xg for 5mins. Supernatant was removed and the cell pellet was resuspended in PBS supplemented with 0.04% BSA and passed through a 40-µm cell strainer. Resulting cell suspensions were either stained with DAPI (BD Biosciences, 1:1000) for dead cell labeling and FACS sorted (AriaFusion, BD Biosciences) or used directly for single-cell RNAseq.

### Single-cell RNA-seq library preparation

Single-cell suspensions were loaded onto the 10x Chromium controller using the 10x Genomics Single Cell 3’ Library Kit v2 (10x Genomics) according to the manufacturer’s instructions. In summary cell and bead, emulsions were generated, followed by reverse transcription, cDNA amplification, fragmentation, and ligation with adaptors followed by sample index PCR. Resulting libraries were quality checked by Qubit and Bioanalyzer, pooled and sequenced using NextSeq500 (Illumina; high-output mode, paired-end 26 × 75 bp). Technical validation (Figure S8) was also performed to show that sorting and fixing of the organoids can be done without affecting the general transcriptomic profile.

### Pre-processing and quality control of scRNAseq data

Raw sequencing data was processed using the CellRanger software (version 3.1.0). Reads were aligned to a custom reference genome created with the reference human genome (GRCh38) and human astrovirus (NC_001943.1). The resulting unique molecular identifier (UMI) count matrices were imported into R (version 3.6.2) and processed with the R package Seurat (version 3.1.3) (Stuart *et al*, 2019) Low-quality cells were removed, based on the following criteria. First, we required a high percentage of mitochondrial gene reads. For the organoid data, all the cells with mitochondrial reads > 10% were excluded. For the tissue data, all the cells with mitochondrial reads > 20% were removed. Second, we limited the acceptable numbers of detected genes. For both types of samples, cells with <600 or >5000 detected genes were discarded. The remaining data were further processed using Seurat. To account for differences in sequencing depth across cells, UMI counts were normalized and scaled using regularized negative binomial regression as part of the package sctransform (Hafemeister & Satija, 2019). Afterward, ileum biopsies and organoids samples were integrated independently to minimize the batch and experimental variability effect. Integration was performed using the canonical correlation analysis (Butler et al, 2018) and mutual nearest neighbor analysis (Stuart et al, 2019; Haghverdi et al, 2018). The resulting corrected counts were used for visualization and clustering downstream analysis and non-integrated counts for any quantitative comparison.

### Clustering and identification of cell type markers

We performed principal component analysis (PCA) using 3000 highly variable genes (based on average expression and dispersion for each gene). The top 30 principal components were used to construct a shared nearest neighbor (SNN) graph and modularity-based clustering using the Louvain algorithm was performed. Finally, Uniform manifold approximation and projection (UMAP) visualization was calculated using 30 neighboring points for the local approximation of the manifold structure. Marker genes for every cell type were identified by comparing the expression of each gene in a given against the rest of the cells using the receiver operating characteristic (ROC) test. To evaluate which genes classify a cell type, the markers were selected as those with the highest classification power defined by the AUC (area under the ROC curve). These markers along with canonical markers for intestinal cells were used to annotate each of the clusters of the ileum samples. For the organoids, The Seurat label transfer routine (Butler et al, 2018) was used to map the cell types from the ileum to the organoid’s cells. Beforehand, immune and stromal cells were filtered and only epithelial cells were used for the ileum reference. Annotation of the organoids cell types was then manually curated using the unsupervised clusters and the identified marker genes. The comparison between the shared cell types from the ileum and the organoids was performed by calculating the Pearson correlation between the averaging normalized gene expression of each cell type.

### Pseudotime inference

To reconstruct possible cell lineages from our single-cell gene expression data. The ileum sample was subset to include only the cell types from con(Street al, 2018)nected linage. The UMAP embedding was then used as input for pseudotime analysis by slingshot (Street et al, 2018). Stem cell was used as a start cluster and Enterocyte and Goblet as the expected end cluster. After pseudotime time was determined and cells were ordered through the resulting principal curves. The genes that significantly change through the pseudotime were determined by fitting a generalized additive model (GAM) for each gene and testing the null hypothesis that all smoother coefficients within the lineage are equal, using the TradeSeq package (Van den Berge et al, 2020). Significantly changing genes were plotted in a heatmap using pheatmap (v. 1.0.10), with each column representing a cell ordered through the pseudotime of either lineage.

### Differential expression analysis

To identify the changes in expression across conditions. Differential expression tests were performed using MAST (Finak et al, 2015), which fits a hurdle model to the expression of each gene, using logistic regression for the gene detection and linear regression for the gene expression level. To reduce the size of the inference problem and avoid cell proportion bias, separate models were fit for each cell lineage and comparisons between mock, 4 hours, and 16 hours post-infection were performed. False discovery rate (FDR) was calculated by the Benjamini-Hochberg method (Benjamini & Hochberg, 1995) and significant genes were set as those with FDR of less than 0.05. Subsequently, Interferon related genes that significantly changed across the conditions (FDR < 0.05) were used to calculate PCA, unsupervised graph-based clustering, UMAP visualization and heatmap. Genes whose mRNAs were found to be differentially expressed were subjected to a gene set overrepresentation analysis using the EnrichR package in R (Kuleshov et al, 2016)

### Multiplex FISH

Organoids were infected, harvested after 16h pi, and embedded in OCT. Section of 10 µm was cut on a cryostat (Leica) and stored at −80 until use. HiPlex (RNAscope) was performed following the manufacturer’s instruction. The RNAscope® HiPlex Assay uses a novel and proprietary method of in situ hybridizations (ISH) to simultaneously visualize up to 12 different RNA targets per cell in samples mounted on slides. Briefly, sections were fixed in 4% paraformaldehyde, dehydrated with 50%, 70%, 100% ethanol, then treated with protease. All the HiPlex probes were hybridized and amplified together. Probes were designed for genes identified as cell type marker and/or corroborated by literature. The probes used were:

| Gene name | Role of the marker | Catalog number | Channel |
| --- | --- | --- | --- |
| SLC2A2 | Mature enterocytes | NPRP-0000689 | T1 |
| SMOC2 | Stem cells | 522921 | T2 |
| MKI67 | Cell cycle | 591771 | T3 |
| CHGB | Enteroendocrine cells | 567681 | T4 |
| FCGBP | Goblet cells | NPRP-0000691 | T5 |
| OLFM4 | Stem-transit amplifying cells | 311041 | T6 |
| BEST4 | Best-M cells | 481501 | T7 |
| FABP6 | Enterocyte lineage | 478701 | T8 |
| LGR5 | Stem cells | 311021 | T9 |
| SPIB | M cells | 551431 | T10 |
| LYZ | Paneth cells | 421441 | T11 |
| HAstVgp1 | Astrovirus | NPR-0001571 | T12 |

The detection was performed iteratively in groups of 4 targets. After washing, cell nuclei were counterstained with DAPI, and samples were mounted using ProLong Gold Antifade Mountant. Imaging was performed with the camera Nikon DS-Qi2 (Nikon Instruments) with the Plan Fluor 40x objective (Nikon Instruments) mounted on the Nikon Ti-E inverted microscope (Nikon Instruments) in bright-field and fluorescence (DAPI, GFP, Cy3, Cy5, and Cy7 channels). The microscope was controlled using the Nikon NIS Elements software. After each round, fluorophores were cleaved and samples moved on to the next round of the fluorophore detection procedures. All images from all rounds of staining were then registered to each other to generate 15 plex images using HiPlex image registration software (ACD Bio). Further brightness and contrast adjustments were performed using Fiji (Schindelin et al, 2012).

## Data and Code Availability

The datasets generated during this study are currently in the process to be submitted to ArrayExpress.

## Supplementary Figures

**Figure S1. Interferons protect human intestinal organoids from HAstV1 infection. A**. Caco-2 cells were infected with HAstV1. At indicated times, the virus was visualized by indirect immunofluorescence for HAstV1 (red). Nuclei were stained with DAPI (blue). **B**. Quantification of the number of HAstV1 infected cells from A. **C**. Same as A, except replication of HAstV1 was assessed by the genome copy number over time using q-RT-PCR. **D**. Same as A, except that the intrinsic innate immune induction of type I (IFNb1) and III (IFNl) IFN were evaluated. **E**. Caco-2 cells were pre-treated for 24h with 2000IU/mL of IFNb1 or 300ng/mL of IFNl1-3. Interferons were maintained during the course of infection and HastV1 infected cells were visualized with indirect immunofluorescence and the number of HAStV1 infected cells was quantified. **F**. Same as E, except replication of HAstV1 was assessed by q-RT-PCR for the genome copy number. **G**. Caco-2 cells, T84 wild-type and T84 cells lacking both the type I and type III IFN receptors (dKO) were infected with HAstV1. 24hpi HAstV1 genome replication was evaluated by q-RT-PCR for the genome copy number. **A-G** Three biological replicates were performed for each experiment. Representative immunofluorescence images are shown. Error bars indicate the standard deviation. Statistics are from unpaired *t*-test.

**Figure S2. General information of single-cell RNA-seq samples from small intestine** Data from. **A-F**. tissue and **G-L**. organoids. Uniform manifold approximation and projection (UMAP) embedding of single-cell RNA-Seq data from human ileum biopsies colored by sample **A**. and **G**., Number of UMI couts **B**. and **H**. and genes **C**. and **I**. Statistics of the single-cell RNA-seq datasets after quality filters are also shown as Violin plot for every sample, Number of cells **D**. and **J**., total UMI counts **E**. and **K**. and detected genes **F**. and **L**.

**Figure S3. Cell type specific features of the small intestine**. Violin plot of the top-ten marker genes for each cell type in the small intestine **A**. biopsies and organoids **B**. features were identified using the receiver operating characteristic (ROC) test. Then selected as those with the highest classification power defined by the AUC (area under the ROC curve). The color in the plots represents the cell type. All genes are reported in tables **S2** and **S3**.

**Figure S4. Comparison of the shared cell types between Ileum biopsies and Ileum derived organoids**. Heatmap showing Pearson correlations between the average expression of every gene across shared cell types between **A**. Ileum biopsies with themself **B**. Organoids with themselves **C**. Ileum biopsies compared to organoids and **D**. Ileum biopsies compared to organoids only using the top 500 variable genes.

**Figure S5. Additional information for Hiplex experiments**. Representative FISH images of all 12 detected genes for cryosection of intestinal organoids. Splitted by hybridization round and colored by the channel used.

**Figure S6. Lineage-Specific Expression Changes in Mock versus 4h and 16h p**.**i**. Volcano plots of genes that are differentially expressed in cells in one time point relative to the other, showing the statistical significance (-log10 adjusted p-value) vs log2 fold change and gene enrichment analysis results for significantly changing genes (FDR <0.05). **A-D**. Enterocyte lineages. **E-H**. Stem cells **I-L**. TA cells **I-P**. Goblet cells **Q-S**. Enteroendocrine cells **T-V**. Best4+ Enterocytes. Selected genes are shown, all genes are reported in Table S4.

**Figure S7. Lineage-specific interferon patterns are conserved across conditions**. UMAP embedding of scRNA-Seq data from human ileum-derived organoids based on the significantly changing ISGs for cells at **A**. Mock, **B**. 4hpi and **C**. 16hpi. **D-F**. Unsupervised clustering of the UMAP data from **A-C. G-I**. The distribution of cell lineages and types in the clusters from **D-F. J-L**. A heatmap of differentially expressed ISGs across the clusters from **D-F**.

**Figure S8. Comparison between live and fixed and FACS sorted and non-sorted single-cell RNA-seq samples from intestinal organoids. A-E**. Fixed and live comparison. **F-J**. Sorted and non-sorted comparison. Boxplots showing the number of UMIs (blue) and genes (red) detected per cell using HIEs dissociated cells on **A**. Live, Methanol, FCS3 and RNAassist Fixed samples. **F**. Samples sorted and non-sorted at mock and 16hp.i **B and G**. UMAP representation colored by sample. **C and H**. UMAP colored by distinct cell clusters, found using unsupervised clustering. **D and I**. Pearson correlation calculated for each gene detected in both samples. **E and J**. Distribution of cells in each cluster according to the sample origin.

