## Supplementary figures and images for "Single-cell transcriptomics reveals immune response of intestinal cell types to viral infection"

### Supplementary Figures

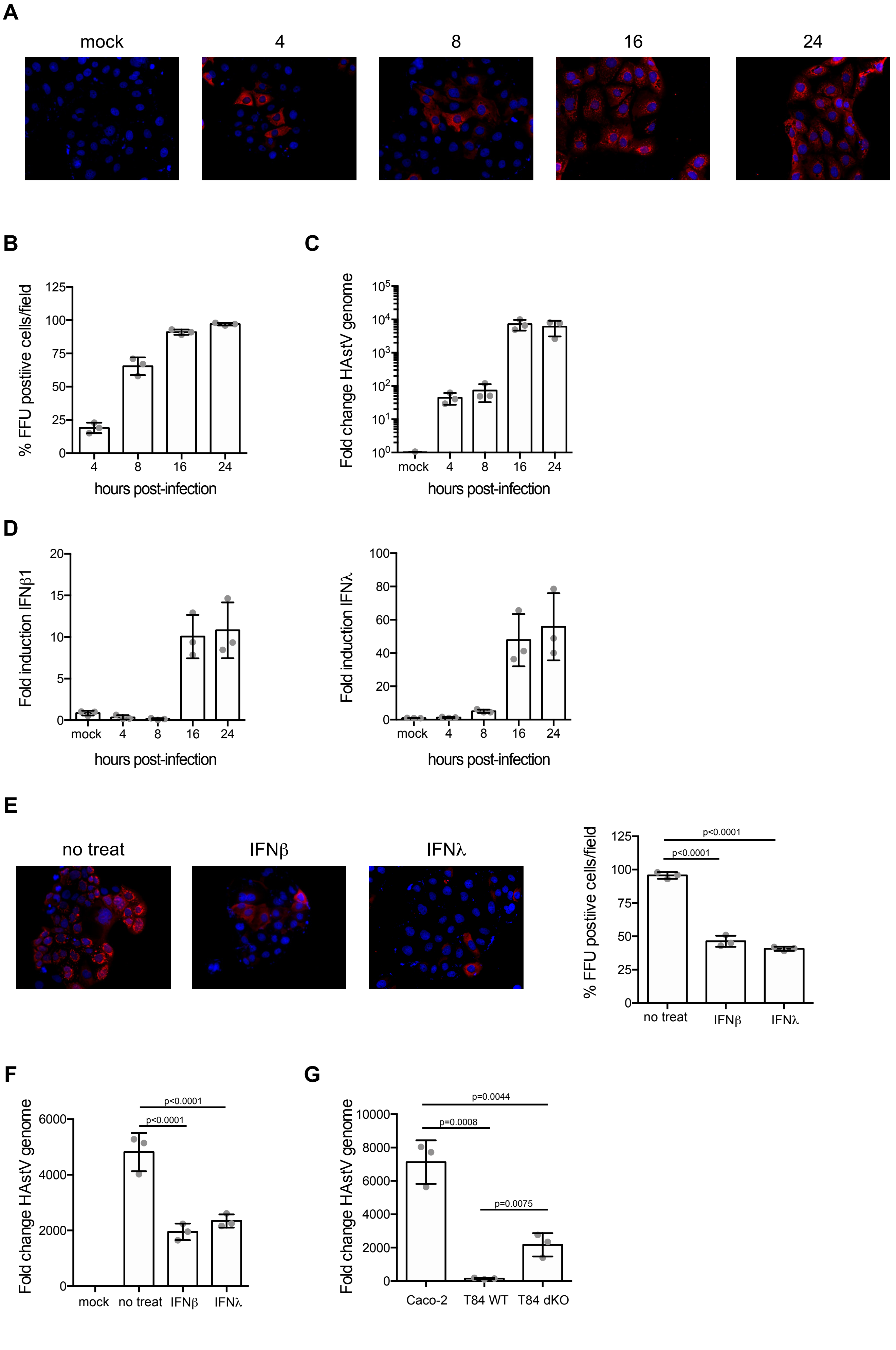

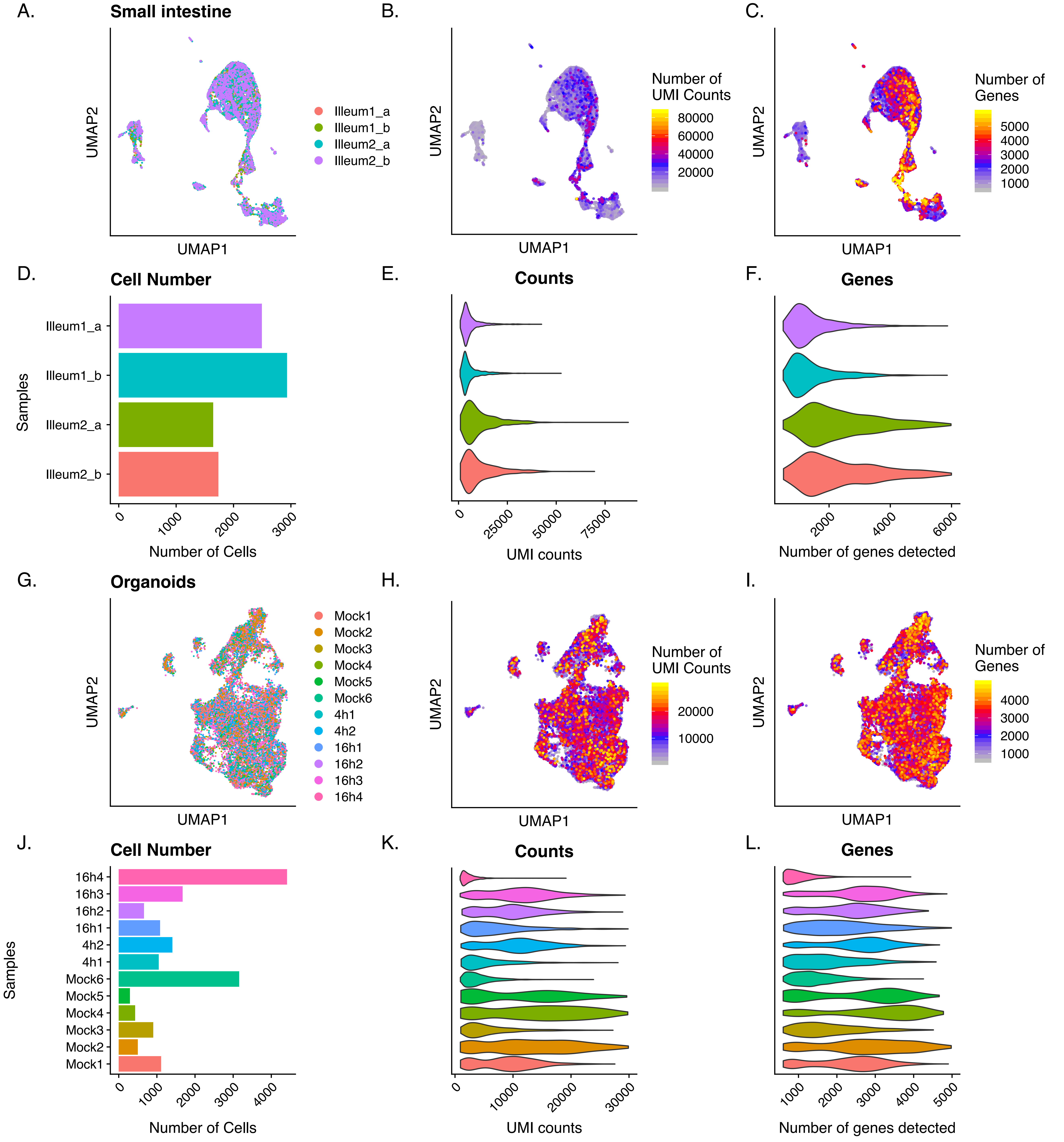

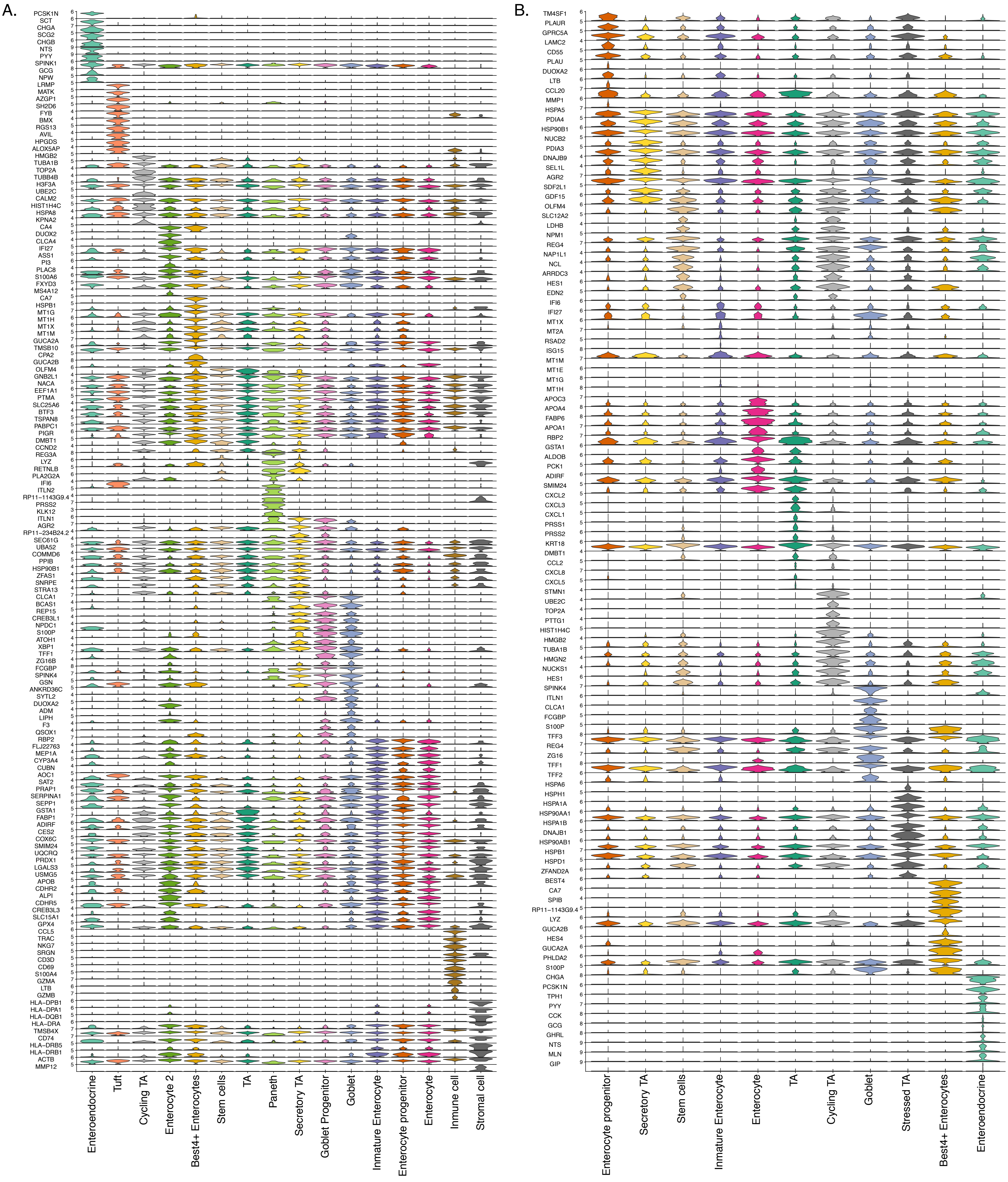

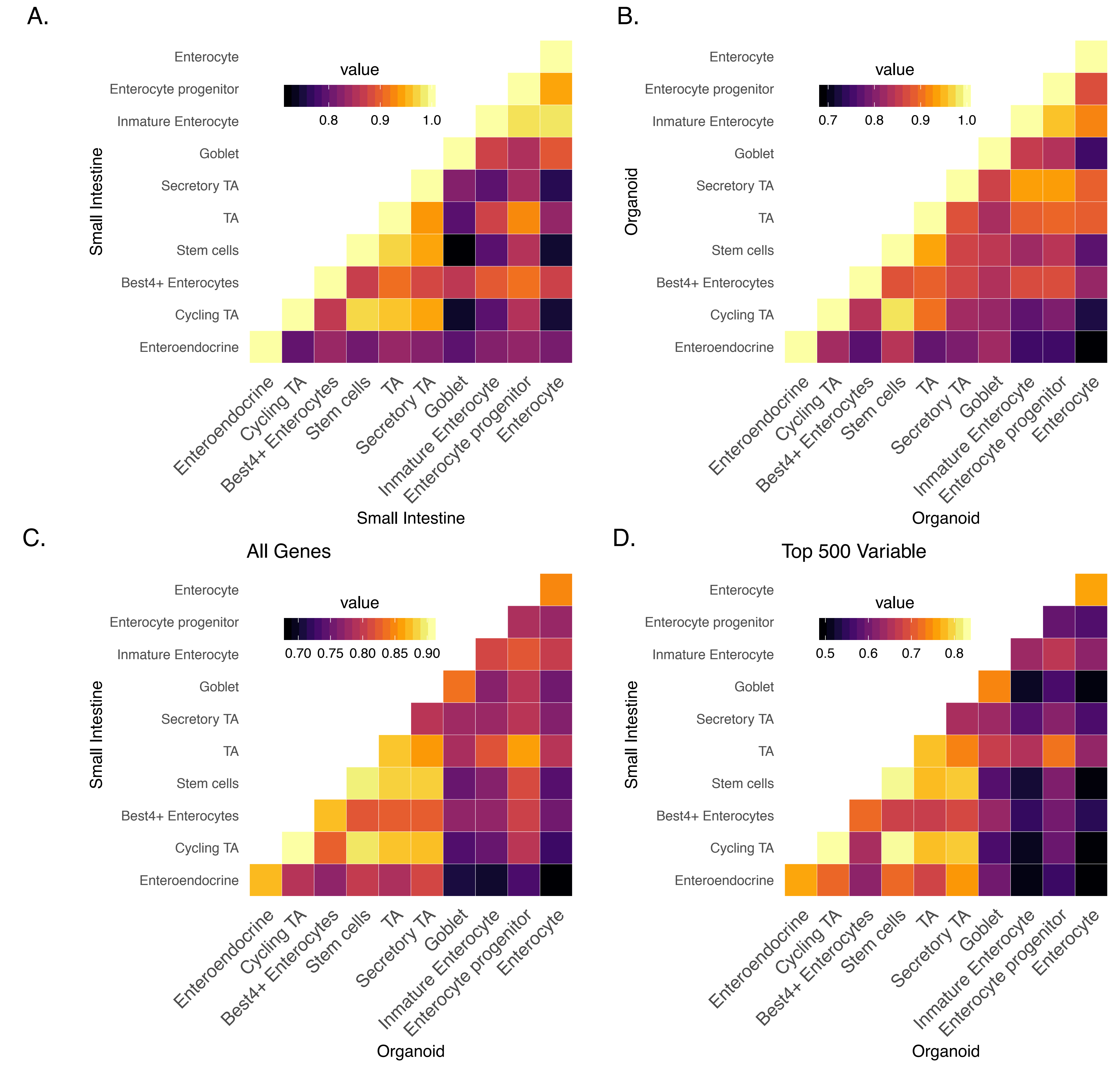

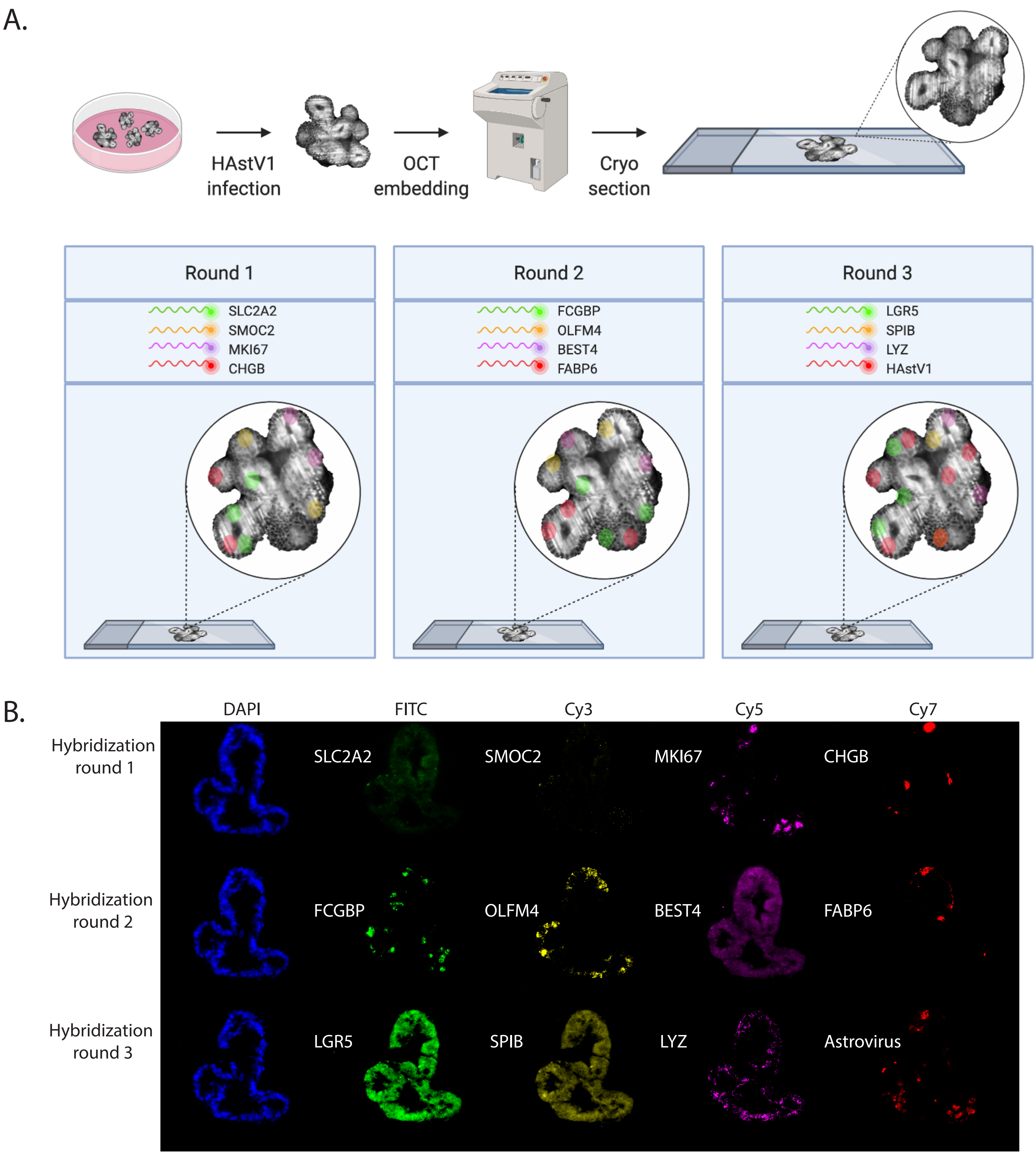

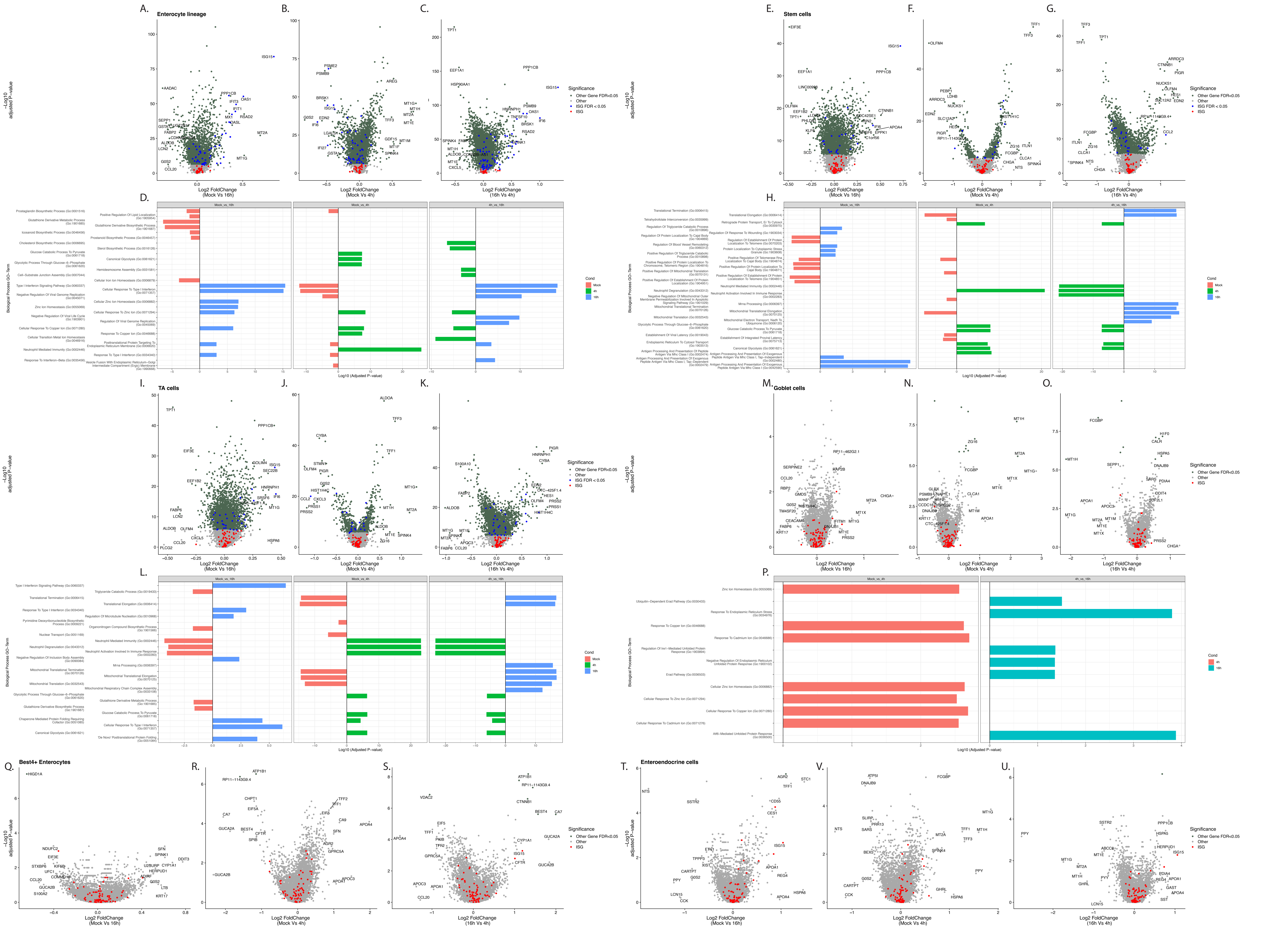

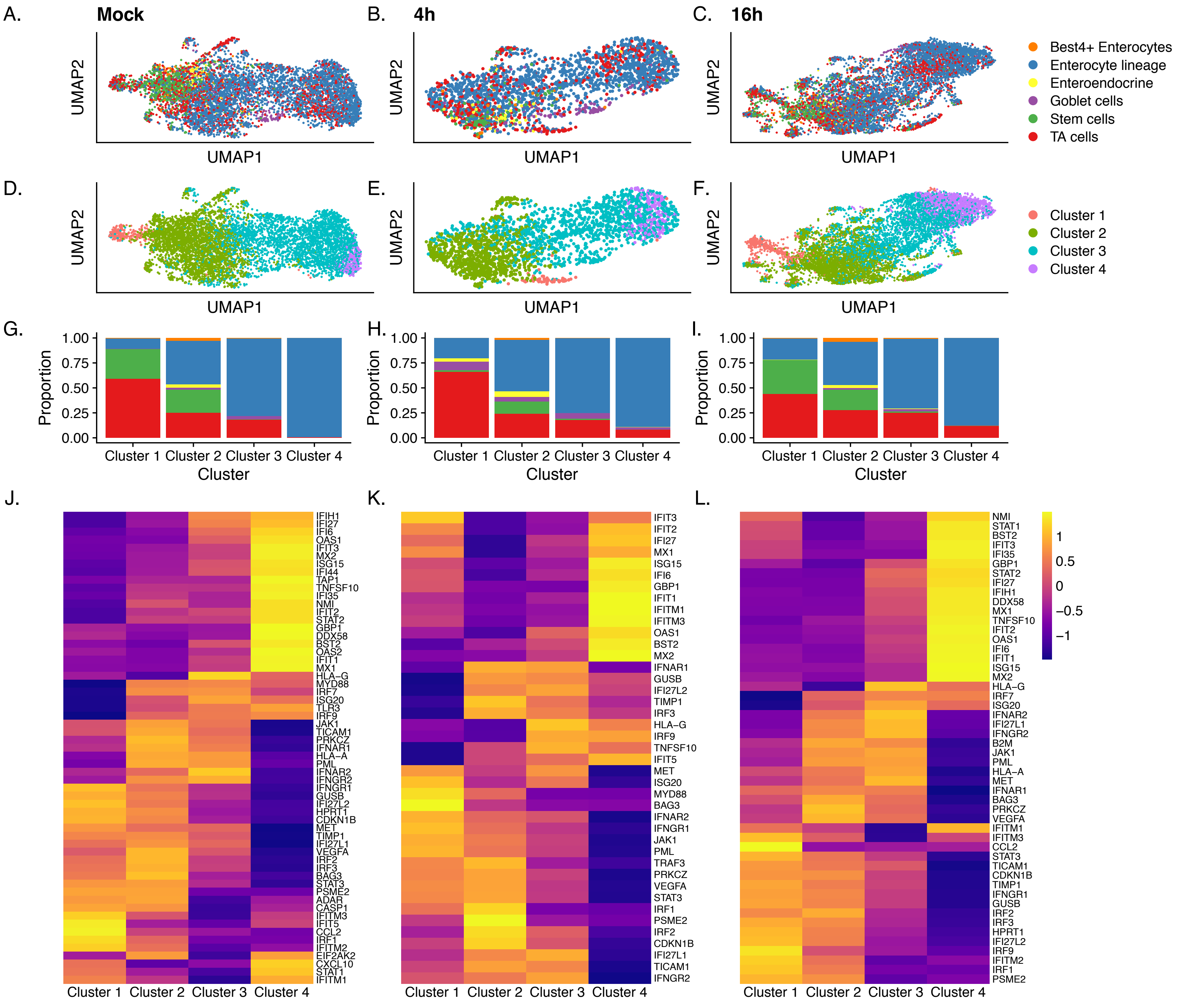

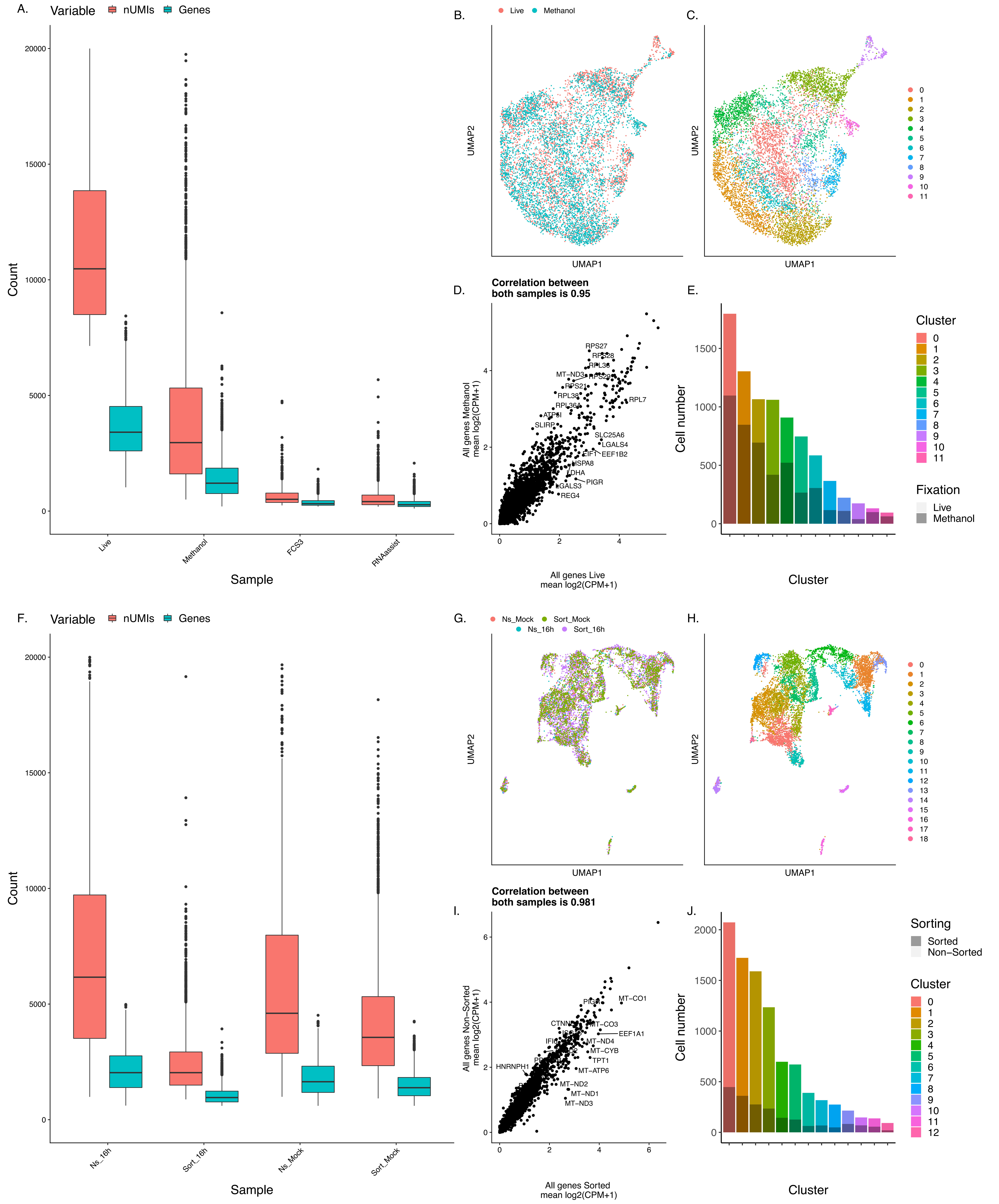
